# Muller’s Ratchet in Action: The Erosion of Sexual Reproduction Genes in Domesticated Cassava (*Manihot esculenta*)

**DOI:** 10.1101/2024.02.14.580345

**Authors:** Evan M Long, Michelle Stitzer, Brandon Monier, Aimee J Schulz, M. Cinta Romay, Kelly R Robbins, Edward S. Buckler

**Affiliations:** Plant Breeding and Genetics Section, School of Integrative Plant Science, Cornell University, Ithaca, NY 14853, USA; Department of Plant Sciences, University of California Davis, Davis, CA, 95616, USA; Institute for Genomic Diversity, Cornell University, Ithaca, NY 14853, USA; United States Department of Agriculture-Agricultural Research Service, Robert W. Holley, Center for Agriculture and Health, Ithaca, NY 14853, USA

**Author notes:** Corresponding Author: Evan Long –.

## Abstract

Centuries of clonal propagation in cassava (*Manihot esculenta*) have engaged Muller’s Ratchet, leading to the accumulation of deleterious mutations due to the absence of sexual recombination. This has resulted in both inbreeding depression affecting yield and a significant decrease in reproductive performance, creating hurdles for contemporary breeding programs. Cassava is a member of the Euphorbiaceae family, including notable species such as rubber tree (Hevea brasiliensis) and poinsettia (Euphorbia pulcherrima). Expanding upon preliminary draft genomes, we annotated 7 long-read genome assemblies and aligned a total of 52 genomes, to analyze selection across the genome and the phylogeny. Through this comparative genomic approach, we identified 48 genes under relaxed selection in cassava. Notably, we discovered an overrepresentation of floral expressed genes, especially focused at six pollen-related genes. Our results indicate that domestication and a transition to clonal propagation has reduced selection pressures on sexually reproductive functions in cassava leading to an accumulation of mutations in pollen-related genes. This relaxed selection and the genome-wide deleterious mutations responsible for inbreeding depression are potential targets for improving cassava breeding, where the generation of new varieties relies on recombining favorable alleles through sexual reproduction.

## Introduction

Cassava (*Manihot esculenta*) is a monoecious root crop grown in tropical regions around the world. Today, cassava is a major caloric source for over 500 million people, with a large number concentrated in Sub-Saharan Africa (Parmar et al., 2017; Ferguson et al., 2019). Cassava is a woody shrub that naturally reproduces through outcrossing facilitated by separate male and female flowers. Although it is naturally perennial, it has been grown as an annual since its domestication 5-10 thousand years ago and vegetatively propagated through stem cuttings (Wang et al., 2014;Parmar et al., 2017). Centuries of selection have generated modern cassava varieties that produce large and abundant roots. This is particularly beneficial in sub-Saharan Africa where it is valued for its ability to grow with minimal inputs in marginally fertile lands, achieving an average of ∼10 tons/hectare fresh root yield (Parmar et al., 2017). With continually rising demands from growing populations and impending difficulties due to climate change and other environmental considerations, breeding efforts for crop improvement in cassava have garnered increasing attention.

One hurdle that impedes contemporary breeding efforts in cassava is high levels of genetic load, made visible through heavy inbreeding depression and low reproductive fitness. Several studies quantified the level of inbreeding depression in self-fertilized cassava, such that a single generation of inbreeding can decrease fresh root yield by >60% (Rojas et al., 2009; de Freitas et al., 2016). These studies likely underestimate the impact of inbreeding depression, as only measured plants that successfully grew from self-fertilized seed can be measured, missing impacts on seed germination and sexual reproduction traits. Further evidence for poor sexual reproductive ability in cassava comes from high variability in flowering time, low numbers of female flowers, high rates of flower abortion, and low seed set (Ceballos et al., 2004; Silva Souza et al., 2020; Oluwasanya et al., 2021). These limitations in seed production and viability limits breeders’ abilities to make the successful crosses inherently necessary for developing new varieties and making genetic gains. Understanding genetic load can help researchers and breeders address these problems hindering genetic improvement of cassava.

One common explanation for genetic load is the accumulation of deleterious mutations in the genome. Domestication has been highlighted as a process that increases fixation of deleterious mutations through linkage with selected loci and a reduction in effective population size (Moyers et al., 2018; Bosse et al., 2019). Cultivated cassava clones have been found to have an abundance of deleterious mutations that are maintained as heterozygous through clonal propagation, likely masking recessive effects (Ramu et al., 2017, Long et al. 2023). The phenomenon known as “Muller’s ratchet” occurs where an absence of recombination, usually through asexual reproduction, allows deleterious mutations to accumulate (Muller, 1964). Clonal propagation and domestication have likely led to the persistence of these deleterious mutations, yet these mutations and their effects have yet to be classified.

By considering its evolutionary history, it is possible to evaluate the deleterious mutations across the cassava genome. Cassava belongs to the Euphorbiaceae, or spurge, family which is a very diverse clade of Malpighiales (Kubitzki, 2014). There are over 8,000 species within the family ranging from tall trees like the rubber tree, *Hevea brasiliensis*, to the ornamental poinsettia, *Euphorbia pulcherrima*. Many species are acclimated to tropical regions, however there are also species that are succulent and adapted to drier regions such as *Euphorbia canariensis*, or canary island spurge. The few common features of uniovulate Euphorbiaceae species are “latex and laticifers, pollen morphology, and ovular and seed coat characters” (Kubitzki, 2014). Cassava is known to have undergone a paleopolyploidy event that is shared with *Hevea brasiliensis* (Pootakham et al., 2017).

In this study we aim to detect selection and genetic load in cassava by capturing evolutionary signals with functional and genomic homology to cassava. Using 27 recently sequenced and assembled Euphorbiaceae species (Long et al. 2023), we perform genetic comparisons across a total of 52 species to detect selection signatures in cassava. We show that evolutionary conservation and accelerated evolution tracks the effects of genetic load through cultivated cassava’s evolutionary history.

## Results

### Genome Assemblies

In conjunction with a breeding study, we sequenced and assembled 27 genomes solely for the purpose of training a machine learning model (Long et al., 2023). In this study, we annotate and characterize these genomes to compare ortholog conservation across the cassava genome. Most of these species were sequenced using short-reads, while 7 species were sequenced using long-read sequencing. In addition to these sampled species, we assembled 11 Euphorbiaceae taxa with publicly available short-read sequences from a botanic garden survey (H. Liu et al., 2019), and collected 15 public reference assemblies (Tuskan et al., 2006; Bredeson, Lyons, Prochnik, Wu, Ha, Edsinger-Gonzales, et al., 2016; Xu et al., 2017; Horvath et al., 2018;Chen et al., 2019; Wei et al., 2020; Wu et al., 2020; R. Zhou et al., 2020; Jalali et al., 2020; J. Liu et al., 2020; Cai et al., 2021; He et al., 2021; B. Li et al., 2021;W. Zhou et al., 2021; Lu et al., 2022) for a total of 53 species (Sup. Table S1). Genome sizes and sequence coverage were estimated through k-mer analysis (Sup. Table S1).

Genome assembly quality was evaluated by assembly size, contiguity, and reconstructed gene space (Fig. 1). The quality of gene-space reconstruction was estimated through Benchmarking Universal Single-Copy Orthologs (BUSCO) and contiguity quality represented by the length of the contig of which 50% of the assembly is contained with that size of contig or larger (N50). These assembled genomes have large variability in contiguity and quality of gene-space reconstruction due to differences in sequencing methods as well as large variability in genome size (Fig. 1). Species assembled from long-reads are of very high quality with N50 and BUSCO values comparable to or higher to many of the previously published reference assemblies.

**Figure 1.**
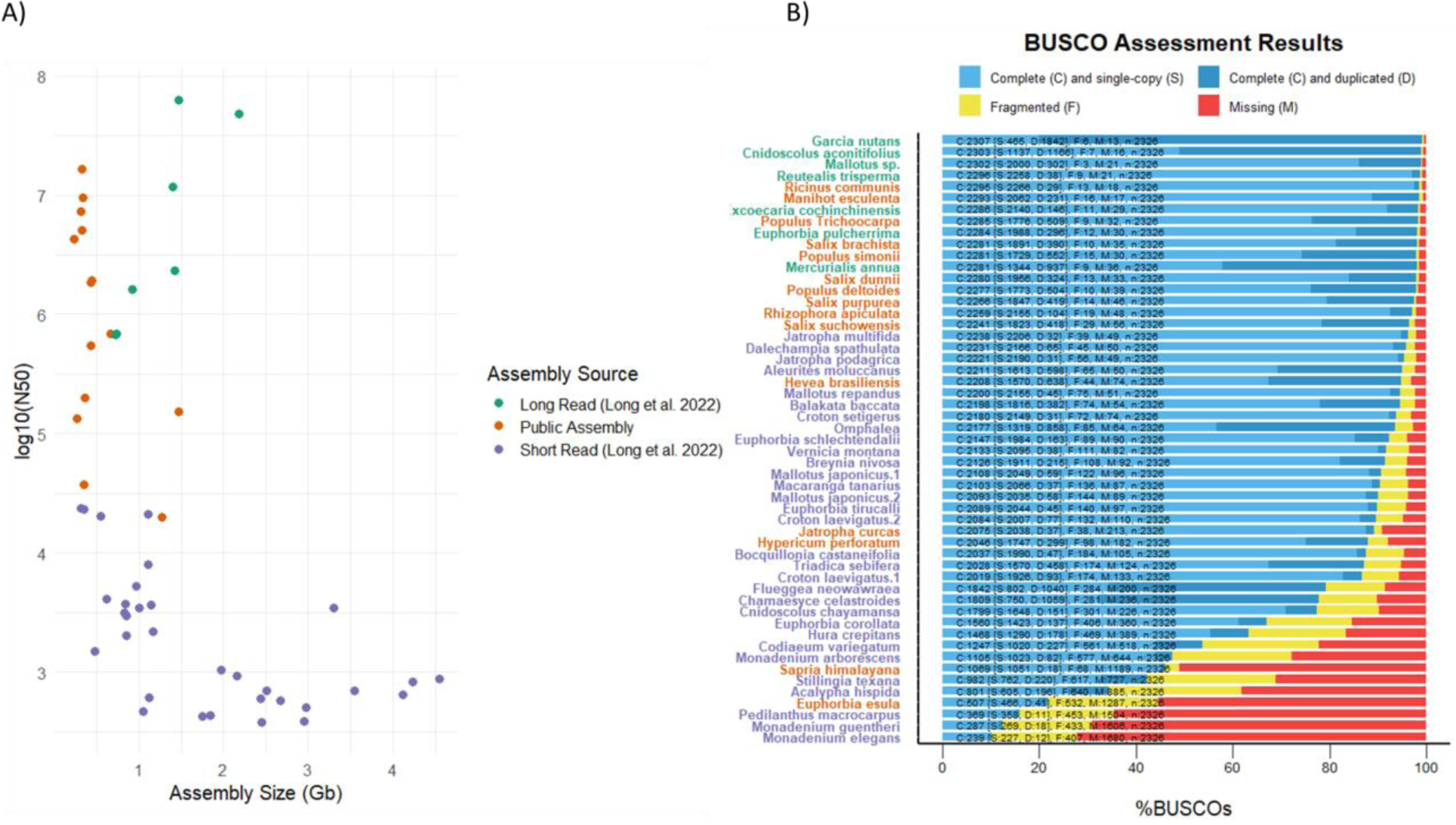
Assembly Quality Statistics. Assembly size and N50 are shown for long and short read assemblies we produced, as well as the public assemblies used in this study (left). Benchmarking Universal Single-Copy Orthologs (BUSCO) scores is plotted with species’ text color matching sources (right).

### Pan-genome Annotations

We combined our long-read assemblies with publicly available Euphorbiaceae genome assemblies to create a Euphorbiaceae gene pan-genome. Our de novo assemblies were annotated using BRAKER2, and genome homology and synteny was produced through the tool GENESPACE (Lovell et al., 2022). This pan-genome defines orthogroups for each gene, including cassava genes. There are over 11k orthogroups that are found in at least 80% of high quality Euphorbiaceae assemblies in our pan-genome (Sup. Fig. S1). These conserved orthogroups account for ∼19k cassava genes (72% of the high-quality genes).

### Evolutionary Conservation Across the Euphorbiaceae Family

We measured the presence of reconstructed gene orthologs across our phylogeny and found a wide distribution of how many genome assemblies had complete homologous sequence for each cassava gene (Fig. 2a). We considered alignments for the single best aligned homologous gene from each genome assembly with at least a 90% aligned length to the cassava homolog. While the distribution of taxa with homologous genes is dependent on assembly quality, there are many genes that are present and completely assembled across a majority of Euphorbiaceae and related species, even from the short-read assemblies. We see very few genes across all 53 species, likely hindered by poor assembly and alignment quality derived from short-read assemblies. The set of cassava genes with very few observations across the Euphorbiaceae may indicate those genes are either unique to cassava, conserved in few species, or may be non-functional annotations thus not conserved.

**Figure 2.**
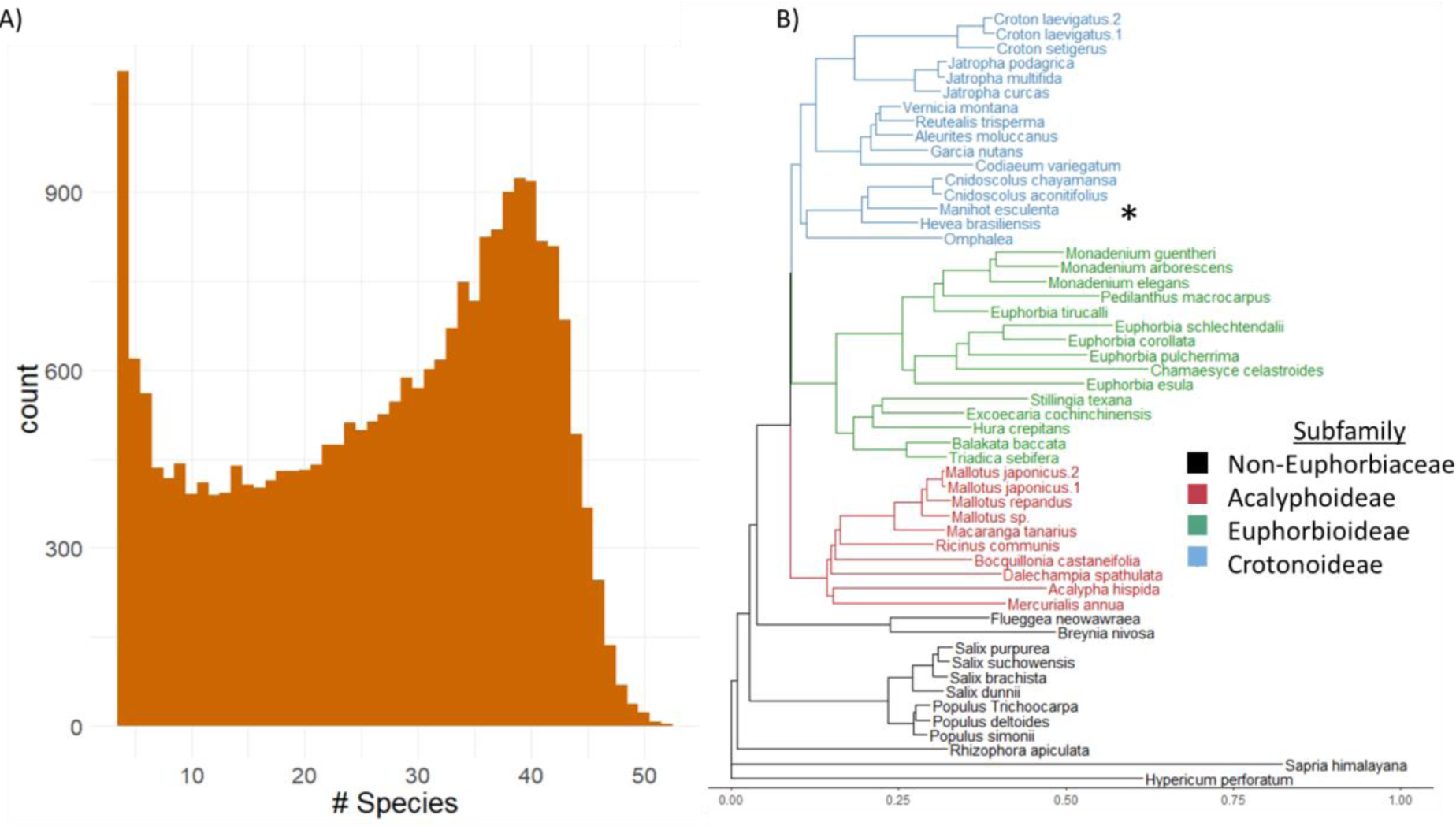
Ortholog occurrence across all assemblies and phylogenetic relationships. An ortholog frequency histogram with the number of species that are represented in each ortholog group across all assemblies (left). Phylogenetic tree created from 4-fold degenerate sites from 1000 randomly selected genes. . The Euphorbiaceae subfamilies are designated by color (right). Cassava (*Manihot esculenta*) is part of the Croronoideae sub-family and designated with “*”.

We constructed a phylogeny to evaluate relationships among these 53 species. A maximum likelihood neutral tree was estimated from 4-fold degenerate sites in a random sample of 1000 genes (Fig. 2b). This phylogeny shows relationships that agree with previously understood taxonomic relationships, including 3 subfamilies: Acalyphoideae, Crotonoideae, and Euphorbioideae (Wurdack et al., 2005). Two species, *Breynia nivosa* and *Flueggea neowawraea*, were previously classified as biovulate subfamily Phyllanthoideae, and are now part of a separate family Phyllanthaceae (Wurdack et al., 2005). Outgroup species, including aspen, willow, and flax, are distantly related but still fall within the order of Malpighiales.

While the contiguity and quality of short-read assemblies is relatively low (Fig. 1A), their assembly of genic regions allowed us to incorporate these species into evolutionary assessment. Many of the short-read assemblies from genomes that were ≤ 1Gbp in size had high contiguity and BUSCO scores, while those from species with larger genome sizes are of lower quality. This is mainly due to common sources of difficulty such as obtaining high enough coverage sequence data and assembling large complex regions with short-read information. Ultimately the utility of these genomes is visible in the amount of reconstructed ortholog space in the cassava genome (Fig. 2A).

### Interspecific Selection Signatures

Using evolutionary conservation and interspecific variation, we analyzed selection signatures across ∼26k cassava gene models that passed quality filters. The measure of dN/dS captures the relative abundance of functional mutations compared to neutral mutations (Kryazhimskiy et al., 2008). We calculated the ratio of nonsynonymous (causing amino acid changes in protein sequence) to synonymous (assumed to be neutral) substitutions across the evolutionary tree (dN/dS) (Fig. 3A). Additionally, we estimated this ratio for substitutions occurring along the cassava branch of the tree. Comparing genome conservation across millions of years allowed us to measure selection on cassava genes. Our results show a large proportion of the cassava genome having a dN/dS < 0.5 implying genes under purifying selection (Fig. 3A). These genes are likely functional and important to be conserved across the Euphorbiaceae family and related species, and to have low tolerance for mutations that disrupt conserved function. Genes with dN/dS ∼1 are likely under neutral selection and are either non-functional or whose function is not currently under selection. Genes with dN/dS >1 are either under positive selection, relaxed purifying selection, pseudogenes, or are poorly estimated due to short gene length (Fig. 3A). For visual clarity we truncated all dN/dS ratios >2, setting their value to 2. We measured relaxation of selection in cassava as the difference between dN/dS of the evolutionary tree and that of the cassava branch. The negative ΔdN/dS indicates a larger dN/dS value in the cassava branch of the tree, or a transition away from purifying selection. By comparing this branch specific ratio to the value estimated across all species we found 167 genes comprising 148 different orthogroups that showed significantly higher dN/dS values along the cassava lineage (Sup. Table S2), suggesting relaxed or positive selection in cassava (Fig. 3B). We found that the reliability of these estimates of dN/dS and significance of separations between cassava and the other species is dependent on length of the coding sequence (CDS) and the number of species with reconstructed orthologs, with a number of short genes experiencing large, but insignificant differences in dN/dS ratios (Fig. 3B&C - lower left quadrants).

**Figure 3.**
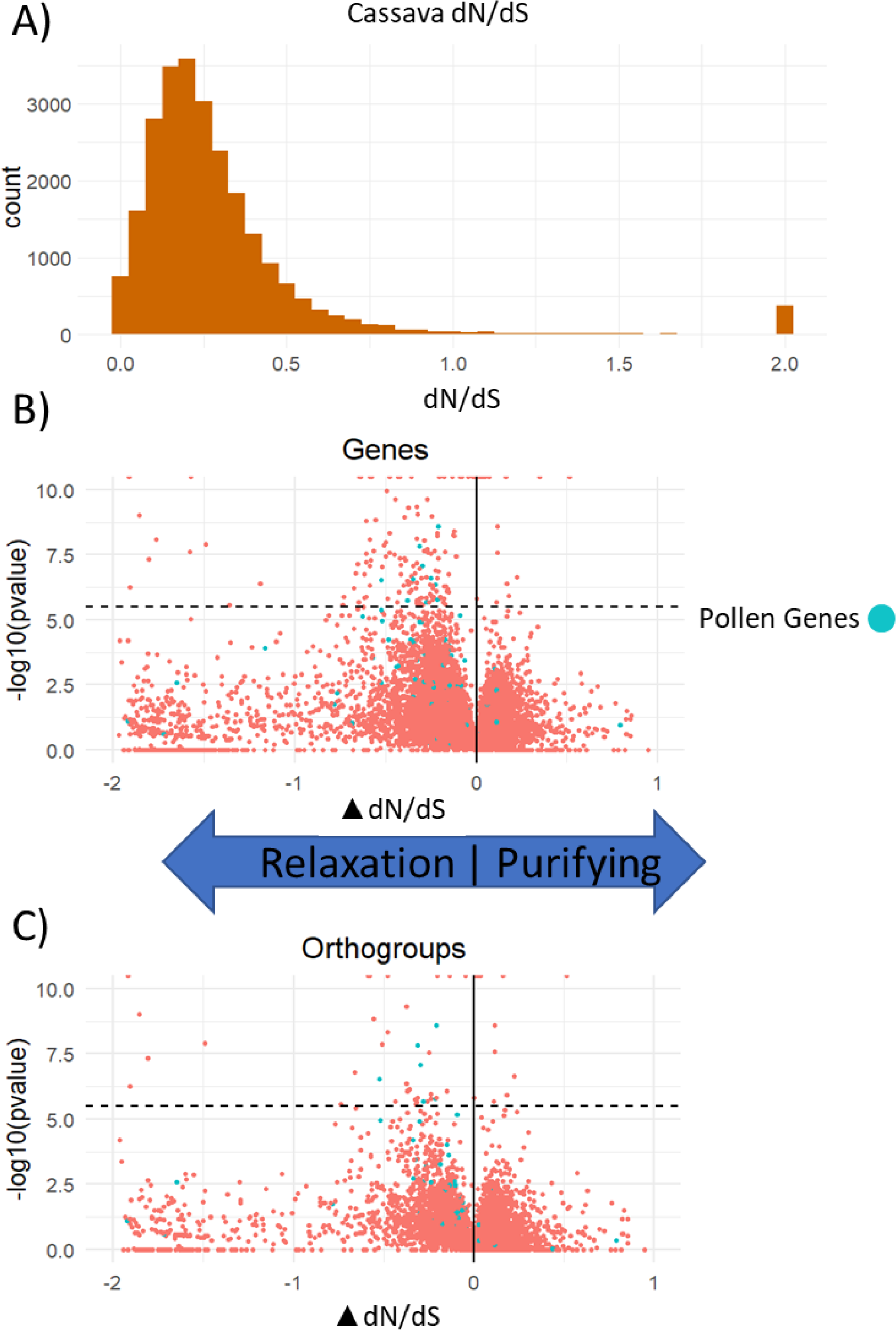
Selection signatures dN/dS gene conservation. A) Histogram of dN/dS values from all genes across cassava, with values greater than 2 plotted at dN/dS=2 (top). B) The difference in dN/dS score between the 52 species used in this study and cassava for each gene in cassava, with the y-axis showing the log ratio test p-value between these two models and dotted line showing multiple test correction significance threshold. C) The difference in dN/dS score between the 52 species used in this study and cassava summarized for each orthogroup in cassava, with the y-axis showing the log ratio test p-value between these two models and dotted line showing multiple test correction significance threshold. Arrows indicate difference in selection in cassava (i.e. ΔdN/dS < 0 implies a relaxation of purifying selection in cassava, or a transition to more positive selection, and ΔdN/dS > 0 implies a stronger purifying selection in cassava)

The ancestor of cassava experienced a whole genome duplication ∼40 million years ago (Bredeson, Lyons, Prochnik, Wu, Ha, Edsinger-Gonzales, et al., 2016), and genes resistant to fractionation have been retained as duplicated genes across the genome. Duplicate genes may show signatures of relaxed selection, if one copy maintains function and the other subfunctionalizes or neofunctionalizes (Flagel & Wendel, 2009). To minimize the possibility of conflating ongoing fractionation with true relaxed selection, we only considered ortholog groups of genes that contained a single cassava gene or where all cassava gene copies passed significance tests, resulting in 48 genes from 47 orthogroups (Fig. 3C).

Since its domestication, selection in cassava has been strongest on root mass. We expected traits unnecessary for clonal reproduction of these large roots to be released from selection. We performed differential gene expression using available RNA-sequencing data from 5 cassava tissues, and found that from among the 48 genes showing relaxed selection, 15 had differentially increased expressed in flowers compared to non-flower tissues (Sup. Fig. S2). Additionally, ∼70% of the *Arabidopsis thaliana* homologs to the 48 genes are most highly expressed in flower, seed, and pollen tissues, (Sup. Table S2).

We then investigated the possible biological functions of this set of genes that show relaxation from purifying selection in cassava compared to the rest of the evolutionary tree. We performed gene ontology (GO) enrichment for GO terms regarding biological processes. The set of 48 genes with significant differences in dN/dS values showed an enrichment for processes involved with pollen and pollen tube development, which exhibited a 20-fold enrichment (Table 1). These ontologies were attributed to six specific genes in cassava: Manes.02G178800, Manes.03G204900, Manes.03G130950, Manes.04G017000, Manes.04G056400, and Manes.08G062900, 4 of which showed differentially higher expression in flowers relative to leaf, stem, fibrous and storage root tissues.

**Table 1.** Enriched GO Terms for Genes Under Relaxed Selection. GO term enrichment produced from “topGO” among genes significantly relaxed from selection (ΔdN/dS <0 and bonferroni multiple test correction significance).

| GO.ID | Term | Annotated | Significant | Expected | Classic Fisher |
| --- | --- | --- | --- | --- | --- |
| GO:0009860 | **pollen tube growth** | 177 | 6 | 0.3 | 2.80E-06 |
| GO:0060321 | **acceptance of pollen** | 12 | 2 | 0.02 | 0.00018 |
| GO:0009409 | response to cold | 652 | 6 | 1.11 | 0.00076 |
| GO:0006893 | Golgi to plasma membrane transport | 27 | 2 | 0.05 | 0.00096 |
| GO:0009651 | response to salt stress | 940 | 7 | 1.6 | 0.00268 |
| GO:0009846 | **pollen germination** | 79 | 2 | 0.13 | 0.00799 |
| GO:0010224 | response to UV-B | 108 | 2 | 0.18 | 0.01452 |
| GO:0015846 | polyamine transport | 10 | 1 | 0.02 | 0.01686 |
| GO:1900039 | positive regulation of cellular response... | 10 | 1 | 0.02 | 0.01686 |
| GO:0032355 | response to estradiol | 10 | 1 | 0.02 | 0.01686 |

### Intraspecific Selection Signatures

Relaxed selection along the cassava lineage identified candidate genes, but it is unclear whether the relaxation of selection occurred across the genus *Manihot,* or after domestication. To determine which pathways were released from selection in domesticated cassava, we used intraspecific data with a sample of 330 sequenced cassava clones. We used two versions of the residual variation intolerance score (RVIS), which uses either excess nonsynonymous (Fig. 4A) or deleterious mutations (Fig 4B) to identify outlier genes. RVIS statistics help control for protein length and differential mutation and drift at the gene level. Nonsynonymous sites were well correlated with the total number of sites (R^2^=0.79). To identify genes with extremes of deleterious variation, we calculated a deleterious RVIS score using variant sites classified as deleterious through evolutionary conservation (DRVIS, Fig. 4B). The regression was much weaker (R^2^=0.14), The genes in the top 5% (n=1287) of RVIS (Table 2) and DRVIS (Table 3) scores, representing those genes with high polymorphism at functional sites, also showed enrichment for pollen related biological processes, among other biological processes.

**Figure 4.**
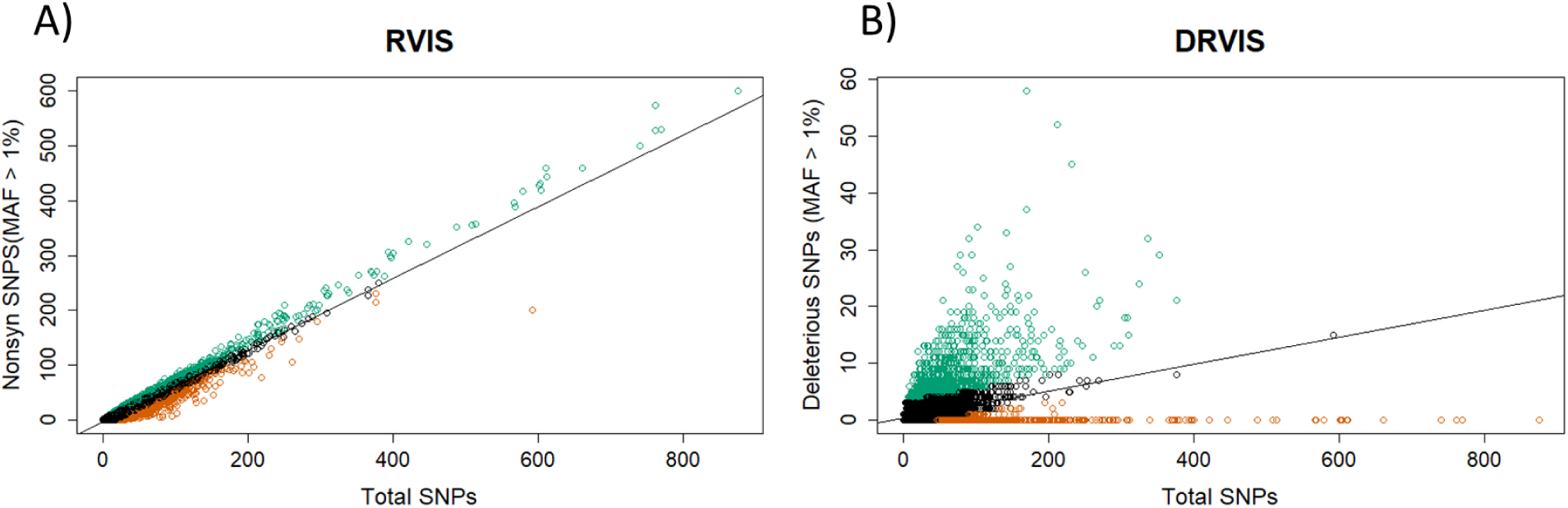
Identification of deleterious mutations in genes using within species Residual Variation Intolerance Scores. Regression for number of nonsynonymous SNPs (A) and putative deleterious SNPs (B) against the total number of SNPs in each gene in cassava. The residuals from each regression give RVIS and DRVIS scores, with the top and bottom 5% (n=1287) of residuals colored as green and orange, respectively.

**Table 2.** Top 10 Enriched Gene Ontology (GO) Terms for Genes with Excessive Non-synonymous mutations (RVIS Genes). GO term enrichment produced from “topGO” among genes in the top 5% of RVIS scores.

| GO.ID | Term | Annotated | Significant | Expected | Classic Fisher |
| --- | --- | --- | --- | --- | --- |
| GO:0006952 | defense response | 3179 | 204 | 114.6 | 1.40E-17 |
| GO:0032922 | circadian regulation of gene expression | 35 | 12 | 1.26 | 1.70E-09 |
| GO:0010483 | **pollen tube reception** | 26 | 10 | 0.94 | 1.10E-08 |
| GO:0030308 | negative regulation of cell growth | 32 | 10 | 1.15 | 1.10E-07 |
| GO:0002239 | response to oomycetes | 142 | 17 | 5.12 | 2.90E-07 |
| GO:0050832 | defense response to fungus | 831 | 56 | 29.96 | 2.10E-05 |
| GO:0010267 | primary ta-siRNA processing | 27 | 7 | 0.97 | 3.60E-05 |
| GO:0009626 | plant-type hypersensitive response | 142 | 16 | 5.12 | 7.30E-05 |
| GO:0009741 | response to brassinosteroid | 158 | 17 | 5.7 | 9.90E-05 |
| GO:0010118 | stomatal movement | 281 | 15 | 10.13 | 0.00015 |

**Table 3.** Top 10 Enriched Gene Ontology (GO) Terms for Genes with a Buildup of Deleterious Mutations (DRVIS Genes). GO term enrichment produced from “topGO” among genes in the top 5% of DRVIS scores.

| GO.ID | Term | Annotated | Significant | Expected | Classic Fisher |
| --- | --- | --- | --- | --- | --- |
| GO:0080168 | abscisic acid transport | 24 | 19 | 1.09 | 9.40E-22 |
| GO:0090332 | stomatal closure | 99 | 20 | 4.51 | 8.70E-15 |
| GO:0006690 | icosanoid metabolic process | 18 | 13 | 0.82 | 2.30E-14 |
| GO:0010496 | intercellular transport | 43 | 15 | 1.96 | 1.90E-13 |
| GO:0015692 | lead ion transport | 19 | 12 | 0.87 | 2.80E-12 |
| GO:0031408 | oxylipin biosynthetic process | 58 | 18 | 2.64 | 4.90E-11 |
| GO:0034440 | lipid oxidation | 79 | 13 | 3.6 | 6.40E-11 |
| GO:0009695 | jasmonic acid biosynthetic process | 63 | 17 | 2.87 | 1.90E-09 |
| GO:0098739 | import across plasma membrane | 79 | 14 | 3.6 | 2.00E-09 |
| GO:0048544 | **recognition of pollen** | 100 | 21 | 4.56 | 3.60E-09 |

Given that pollen related traits are enriched among the extremes of both interspecific and intraspecific measures of genetic load, we further investigated all 348 genes that had pollen related GO terms, irrespective of whether they reached significance in individual tests. We performed Chi-square tests for significant differences in distributions of ΔdN/dS, RVIS, and DRVIS between 348 pollen related genes and all other cassava genes (Fig. 5). We found ΔdN/dS (p-value=0.027) and RVIS (p-value=0.0055) to be significantly lower in pollen related genes than all other genes, while DRVIS (p-value=3.3e-05) was significantly higher. All these effects, while significant, are small, but are consistent with relaxation from selection among pollen genes in cassava.

**Figure 5.**
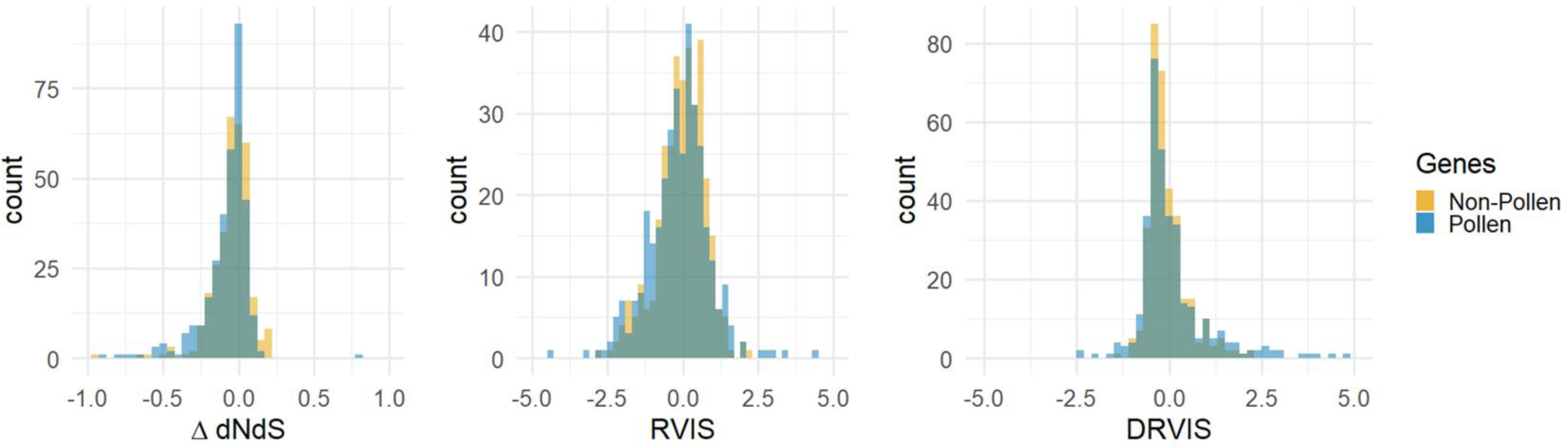
Distributions of ΔdN/dS, RVIS, and DRVIS between pollen and non-pollen related genes. Histograms are shown between pollen (blue) and non-pollen (orange) related genes for ΔdN/dS, RVIS, and DRVIS. Non-pollen related genes are subsampled to an equal number of genes for visual comparison.

In addition to dN/dS, RVIS, and DRVIS we also used the McDonald–Kreitman (MK) test to evaluate selection measures. The MK test uses a combination of intraspecific and interspecific functional variation to measure selection with a positive value (α>0) indicating the proportion of substitutions fixed by positive selection. GO term enrichment was performed on genes in the top 5% of the MK test α value (α=1), and found many functions related to plant defense, like RVIS (Table S3). The MK test α value also showed enrichment for pollen tube functions, though overall no significant difference across all pollen genes (Sup. Fig. S2). Negative estimates of alpha are common in plant populations (Gossmann et al., 2010), as we observe for the majority of genes in cassava. This may be due to low effective population sizes in cassava, intensified by the population contractions seen during the domestication and improvement bottlenecks (Ramu), altering the landscape of segregating variation..

### Chromosome Evolution

Knowing that the cassava genome has experienced a paleotetraploidy event, we examined previously characterized homeologous chromosomes to look for any asymmetrical measures of conservation and selection. We took the average genetic distance to each other genome assembly across all cassava genes. These distances were then compared across homeologous chromosomes to look for any evidence of chromosome lineage resulting from an allopolyploid event (Sup. Fig.S4). We found that with the current resolution provided from the genome assemblies in this study, there is no evidence for allopolyploidization, however this cannot be certain from the available information.

We also examined our selection metrics across the cassava chromosomes to look for any signatures of biased fractionation, due to one homeologous chromosome being selectively conserved over the other (Sup. Fig. S5). While regions of some chromosomes show some signatures of decay (low gene densities, low recombination rates, high dN/dS values), there does not appear to be any evidence for whole chromosome degradation. However, among genes with multiple paralogues, there is evidence for gene sub-functionalization or degradation as seen by one gene copy showing high conservation over other gene copies (Sup. Fig. S6).

## 1. Discussion

The intricate interplay between natural selection, domestication, and prolonged clonal propagation has shaped the cassava genome over the course of its cultivated history. Using the evolutionary principle of Muller’s Ratchet as a framework, we test the hypothesis that clonal propagation after domestication, and the absence of recombination, has led to the accumulation of deleterious load in the cassava genome. Employing interspecific and intraspecific analyses, we evaluate selection signatures across the genome, corroborate with tissue specific expression, and investigate affected gene pathways, and find evidence for relaxed selection in pollen and flower related genes.

### Evolutionary Selection Signals

Surveying millions of years of evolution by sampling taxa across the Euphorbiaceae, we measured selection on each cassava gene. Most genes under differential selection are relaxed along the cassava lineage (Fig. 3B&C), and are enriched for pollen related functions. A loss of pollen function would be catastrophic for a wild plant, but domesticated cassava is propagated clonally. Asexual reproduction may allow the accumulation of mutational load in sexual function genes. Related breakdowns in genes involved in pollen function have been observed between outcrossing wild species and clonally propagated cultivars, for example in potato (Hardigan et al., 2017) and the ornamental *Ranunculus* genus (Kocot et al., 2022). Even wild species may show signatures of this process. In the tree species *Populus tremuloides,* male fertility declines with the clonal age of an individual, potentially due to the accumulation of somatic mutations (Ally et al., 2010). In *Decadon verticillatus*, a transition to asexual reproduction led to the loss of sexual compatibility, primarily through pollen dysfunction (Eckert et al., 1999).

Comparative evolutionary signal supports a role in sexual reproduction of the 48 relaxed cassava genes. Two of these genes, Manes.02G178800 and Manes.08G062900 are annotated as exocyst and secretory complexes that have been experimentally shown to be necessary for proper pollen development in *Arabidopsis thaliana* (Marković et al., 2020). Two more genes, Manes.03G130950 and Manes.04G017000, are annotated as members of the ATP-binding cassette transporter family, whose homologs are essential for anther and pollen exine development in rice (Qin et al., 2013).

### Intraspecific Selection Signals

Genes under relaxed selection in the reference genome of *Manihot esculenta* relative to other taxa could represent episodes of selection that happened during speciation, or during domestication. To disentangle these effects, we used genotyped cassava clones to provide a within species perspective on selection. Similar to the evolutionary signal seen in comparative genomics across the Euphorbiaceae, genes with pollen related functions showed enrichment for functional variation, measured by RVIS and DRVIS, between cassava clones. Further analysis of RNA expression data in cassava supports the sexual reproduction related functions of the 48 genes that showed relaxed selection, with many of them showing differentially increased expression in flower tissues (Sup Fig. S2).

Other significantly enriched GO terms from these population level analyses included defense response functions. This may result from disruptive positive selection, as high amounts of functional variation can be beneficial. Selection for high diversity of plant defense genes has been previously shown in plants (Zhang et al., 2014; Zheng et al., 2016), and these defense response genes may be relevant for common diseases afflicting cultivated cassava such as cassava mosaic virus or cassava brown streak disease. Genes with functions related to circadian rhythms also showed evidence for positive selection, which agrees with previous understanding of gene functions in plants commonly under positive selection (Michael et al., 2003). While gene ontology terms are a coarse estimation for function, these and the other significant functional elements may be further investigated to shed light on molecular functions behind cassava evolution, domestication, and cultivation.

Sexual recombination and seed production are essential to combine favorable alleles to create improved cultivated varieties. Low fruit and seed production hinders breeding efforts in cassava. Low rates of female flower production and large variation in flowering times have been targets to alter for increased cassava seed production (Oluwasanya et al., 2021). Flowering induction is only part of the problem, however, as many studies have reported flower abortion rates of over 80% (Sousa et al., 2021; Ramos Abril et al., 2019; Ukwu et al., n.d.). Studies have also found genotypic variability in pollen viability in cassava, and that self-incompatibility does not explain this variability (Sousa et al., 2021; Ramos Abril et al., 2019). It has been suggested that fruit abortion in out-crossing species may be due to deleterious mutations (Wiens et al., 1987). Multiple studies have shown low pollen amounts and low pollination rates in cultivated cassava crosses compared to wild progenitors and other Manihot species (da Silva et al., 2018; Jennings, n.d.; Vieira et al., 2012). Previous studies on the relationship between cassava clonality on deleterious mutations have shown an unexpected lack of correlation between recombination and deleterious mutations (Ramu et al. 2017), supporting the conclusion that these mutations are indeed being enriched through absent recombination from clonal reproduction.

## Conclusion

This work has produced a deep evolutionary resource for the evaluation of selection and deleterious mutations in cassava. Evolutionary conservation across the Euphorbiaceae family can help determine the functional importance of genes across the cassava genome. Following Muller’s ratchet, the lack of sexual reproduction and recombination has led to deleterious effects on sexual viability in cassava. Understanding the impacts of clonal propagation and the genetic load in genes related to sexual reproduction can help overcome the reproductive hurdles in cassava breeding. These results address only one aspect of genetic load and deleterious mutations in cassava, but the evolutionary resource produced has the potential to address many more in the future.

## Methods

### Sequencing and assembly

We gathered a total of 52 related species in addition to cassava, 27 of which we sequenced and assembled, to evaluate evolutionary conservation and selection across the cassava genome. In order to maximize the amount of evolutionary time sampled, while maintaining reliable alignments to cassava, we sampled 26 species across the Euphorbiaceae family, to which cassava belongs. These species were collected from : the Germplasm Resources Information Network and contributions from many botanic gardens across the United States including: Denver Botanic Garden, the Missouri Botanic Garden, the Montgomery Botanic Garden, the National Botanic Garden, the National Tropical Botanic Garden, The New York Botanic Garden, and the US Botanic Garden.

We then extracted DNA from leaf tissue and sequenced these individuals using Illumina NovaSeq-6000. Genome sizes were estimated using k-mer spectra created using ‘jellyfish’ (Marçais & Kingsford, 2011) with a k-value of 21, in order to estimate sequence input coverage for assembly (https://bioinformatics.uconn.edu/genome-size-estimation-tutorial/). Additional short-read sequences were downloaded from SRA (https://www.ncbi.nlm.nih.gov/sra/) corresponding to 11 unspecified Euphorbiaceae taxa that were previously part of an effort to digitize a botanic garden (H. Liu et al., 2019). We then used a short-read sequence assembler MEGAHIT (D. Li et al., n.d.), with modified parameters of “-m 0.2 --no-mercy --min-count 3 --k-min 31 --k-step 20” to create contig assemblies. These parameters follow recommendations for genome assemblies of complex genomes with >30X sequence coverage (https://github.com/voutcn/megahit).

We additionally obtained long-read sequences using PacBio Sequel II for 7 species among our sampled Euphorbiaceae taxa representing a diverse sample across the family. These include: *Cnidoscolus aconitifolius*, *Euphorbia pulcherrima* (poinsettia), *Excoecaria cochinchinensis*, *Garcia nutans*, *Mallotus sp.*, *Mercurialis annua*, and *Reutealis trisperma*. These sequences were assembled using Hifiasm (Cheng et al., 2021) utilizing default settings. An additional 14 genome assemblies from other related species were downloaded from SRA (https://www.ncbi.nlm.nih.gov/sra/) and added to our assembled genomes resulting in a total of 52 species, excluding cassava (Sup. Table S1).

Genome quality metrics were calculated to inform their usefulness in later analyses. We calculated and reported assembly size and the length of the shortest contig for which longer and equal length contigs cover at least 50% of the assembly (N50). Benchmarking Universal Single-Copy Orthologs (BUSCO) analysis was performed using the eudicot ortholog lineage database (Simão et al., 2015). This metric gives a rough estimate of how well the gene-space is captured by the assembled genome, while also giving a snapshot of the level of gene duplication.

### Pan-genome Annotation

For the long read assemblies, we performed genome annotation using the BRAKER2 protein homology pipeline (Brůna et al., 2021). This BRAKER2 pipeline utilizes ProtHint (Brůna et al., 2020) and a protein database consisting of the Viridiplantae clade to produce de novo gene annotations. We combined these assembled genomes with other Euphorbiaceae public assemblies of *Manihot esculenta*, *Hevea brasiliensis*, *Ricinus communis* to create a pan-genome panel. Orthogroup and synteny analyses were performed using GENESPACE (Lovell et al., 2022) using the default pipeline. These orthogroups were used to define homologous genes across cassava for all analyses.

### Multiple Sequence Alignment

To compensate for the large variation in assembly quality, we used a limited alignment process to align fully reconstructed genes in each species. Cassava transcripts were aligned to each genome while tracking UTR, intron, and exon positions. Exonic regions in the target genomes that could be aligned by ≥ 90% of the transcript and had the highest alignment score consolidated with the query transcript. Multiple sequence alignments were created using MAFFT ‘--ep 0 --genafpair --maxiterate 1000’ for each cassava transcript. While this methodology ignores duplicated copies of genes in target genomes, it simplifies analyses by avoiding errors introduced from fragmented assemblies and polyploidy. Additionally, to enable in-frame protein coding analysis of across homologous genes, any positions with gaps in the cassava transcript were removed from the multiple sequence alignment.

### Gene Tree Analysis

We performed phylogenetic analyses to assess gene evolution and selection signatures.

First, we generated maximum likelihood trees using RAxML ‘-m GTRGAMMA -p 12345’ (Stamatakis, 2014) for every transcript with a minimum of four aligned genomes. To estimate the neutral evolutionary tree of these species we randomly sampled 1000 genes and concatenated the 4-fold degenerate sites (sites where mutations produce no amino acid changes) from their multiple sequence alignments. We rooted this tree to the Malpighiales species *Hypericum perforatum*.

Next, we used the Phylogenetic Analysis by Maximum Likelihood (PAML) suite of tools to evaluate gene and site level conservation (Yang et al 2007). For protein coding analysis we executed two different models of the PAML codeml tool. The models differ by either treating the tree as having one ω ratio parameter for the entire tree or allowing two ω parameters: one for the cassava branch and one for the rest of the tree. These models report the ratio of nonsynonymous to synonymous variants (dN/dS) across the tree as well as the likelihood of the given model. To aid in interpretability of dN/dS >>1, we set the maximum dN/dS=2, thresholding all values greater than this to 2. We interpreted the difference between these models as a method of detecting a difference in selection between cassava and the rest of the tree and performed likelihood ratio tests to verify the significance of any differences between these models. Additionally, we performed PAML baseml analysis for each transcript multiple sequence alignment giving an evolutionary rate for every base-pair position. For the subset of pollen-related genes found to be under significant relaxed or positive selection, we analyzed protein evolutions using a PAML Branch-Site model. This model is used to identify which amino acid substitutions are responsible for the differences between our target branch, Cassava, and all the other species.

### Gene Model Selection

We used our evolutionary information to filter down the ∼64k and ∼33k transcript and gene from the CassavaV7.1 annotations, respectively, to 26k gene models with a single most conserved transcript. First, we filtered out any transcripts that had no annotated untranslated regions, as this suggests it has minimal RNA evidence. Next, we retained gene models that were found in at least one of the other 52 assembled species, as de novo gene generation without any homology at this timescale is likely rare. The best transcript for the remaining genes was decided by looking at the lower dN/dS ratio indicating the most evolutionary conservation and potential for functionality. Any remaining multiple transcripts at a gene were filtered to the longest transcript, which, while not necessarily ideal, provides us with a single transcript for future analysis for each of the ∼26k genes.

### Intraspecific Selection Signatures

We used a large panel of cassava clones to assess intraspecific measures of selection. From the haplotype map found on cassavabase.org (Fernandez-Pozo et al., 2015), we filtered down to 330 cassava clones with at least 10X average genome coverage. We then filtered variant sites to biallelic single nucleotide polymorphisms (SNPs) with less than 20% missingness and minor alleles having at least 3 occurrences. We calculated the Residual Variation Intolerance Score (RVIS) (Petrovski et al., 2013), by regressing the number of nonsynonymous SNPs with a minor allele frequency ≥ 1% onto the total number of SNPs in each gene, then scoring each gene’s studentized residual from this regression. To further focus on those functional variations likely to represent deleterious load, we performed the same analysis on nonsynonymous sites flagged as putatively deleterious. These deleterious mutations were classified previously (Long et al., 2023) and are sites with a baseml evolutionary rate < 0.5, a Sorting Intolerant From Tolerant (SIFT) (Ng & Henikoff, 2003) score ≤ 0.05, and a minor allele frequency < 20%. The RVIS methodology was then repeated to form a score we term here as deleterious RVIS (DRVIS).

We also combined interspecific and intraspecific measures of functional variations to test for positive selection. The test performed was the “McDonald–Kreitman test” (McDonald & Kreitman, 1991) which can be calculated as α=1-((pN/pS)/(dN/dS)), where pN/pS is the ratio of nonsynonymous mutation sites to synonymous mutation sites within a population. We used the cassava hapMap (Ramu et al., 2017) population to determine pN/pS, using a minor allele frequency cutoff of 10%, and calculated α. Similar enrichment tests to those previously described were performed with α, where the alpha value (α>0) describes the proportion of sites fixed by positive selection (Sup Table S3, Sup Fig. S3).

### Selection Evaluation

We collected gene ontologies (GO) through homology to the TAIR10 Arabidopsis thaliana genes (https://www.arabidopsis.org/). BLASTP was performed between CassavaV7 and TAIR10 to determine homologous genes. GO term enrichment was performed using the ‘topGO’ package in R, analyzing GO terms for biological processes in regard to each of our selection measures (Tables 1-3, Sup Table S3). We used public databases of Arabidopsis thaliana gene expression atlases (Waese et al. 2017, https://bar.utoronto.ca/eplant/) to compare tissue specific expression of orthologous genes to the set of 48 genes with relaxed selection signatures (Sup Table S2) (Scmid et al. 2005;Nakabayashi et al. 2005).

Cassava underwent a paleopolyploidy event and many genes contain duplicates throughout its genome. Because of this genome duplication, we addressed our measures of selection on individual genes as well as consolidating their metrics by ortholog group, by recording the least extreme value (i.e. smallest absolute value or least significant p-value). Additionally, we analyzed differences in selection signatures between paralogues within ortholog groups (Sup. Fig. S6) and measured evolutionary distances to each assembled species and known homeologous chromosome pairs (Bredeson, Lyons, Prochnik, Wu, Ha, Edsinger-gonzales, et al., 2016) to determine if there is any evidence for allopolyploidization, which would show asymmetric similarity to a relative of a putative diploid progenitor (Sup Fig. S4). Additionally, we binned our selection measurements (dN/dS, RVIS, DRVIS, etc) and other genome annotations (gene density, genetic map, and domestication sweeps (Ramu et al., 2017)) into 250 kb bins to examine large scale signatures of asymmetric selection across homeolgous chromosome (Sup Fig. S5)

### Differential Expression

Differential expression between flower (mixed female and male inflorescences) and non-flower tissues was performed for the 48 genes found to be under relaxed selection. RNA sequence counts across 5 different tissues and 150 cassava clones from a previous study were used to evaluate gene expression (Ogbonna et al., 2021). Differential expression analysis was performed using R package “DEseq2” (Love et al. 2014), with flower tissue expression compared . Log2Fold changes in expression with significance levels were reported (Sup. Fig. S1).

## Data Availability

The inputs to analyses, code to reproduce tables and plots, and summary tables can all be found on the github repository (https://github.com/em255/CassavaEuphorbiaceaeGeneEvolution).

## Acknowledgements

We would like to acknowledge the many germplasm sources that contributed tissue for sequencing (Sup. Table S1) including: the Denver Botanic Garden, Germplasm Resources Information Network, the Missouri Botanic Garden, the Montgomery Botanic Garden, the National Botanic Garden, the National Tropical Botanic Garden, The New York Botanic Garden, and the US Botanic Garden. Their support was essential in sampling the vast number of species used in this study. We would also like to thank the NextGen Cassava community for the contribution of cassava genotype data that we used from cassavabase.org.

