## Supplemental Tables + Figures for "Muller’s Ratchet in Action: The Erosion of Sexual Reproduction Genes in Domesticated Cassava (*Manihot esculenta*)"

### Supplemental Figures

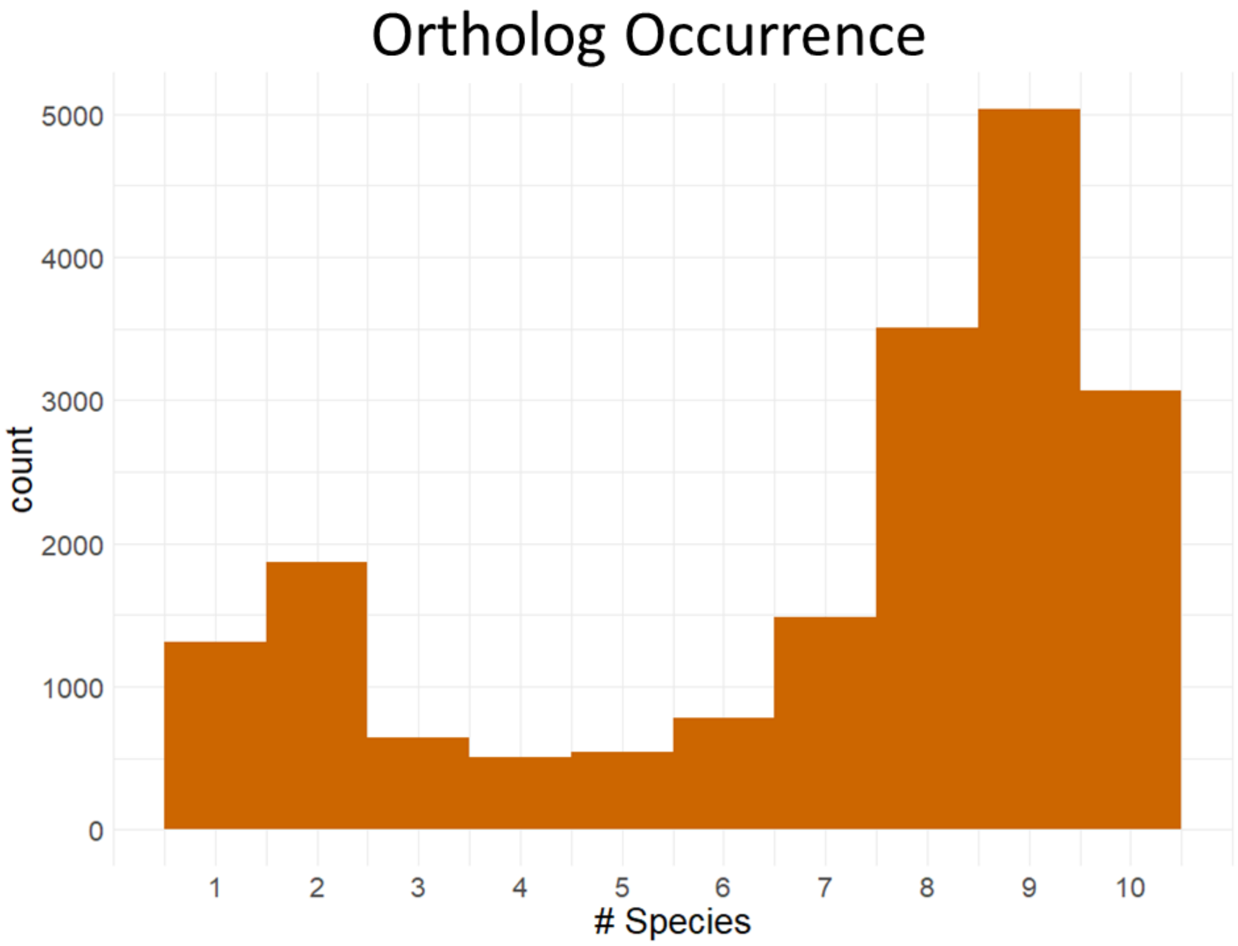

Supplemental Figure S1. Ortholog occurrence across high quality assemblies. An ortholog frequency histogram with the number of species, from among the 10 high quality genomes used in GENESPACE, that are represented in each ortholog group.

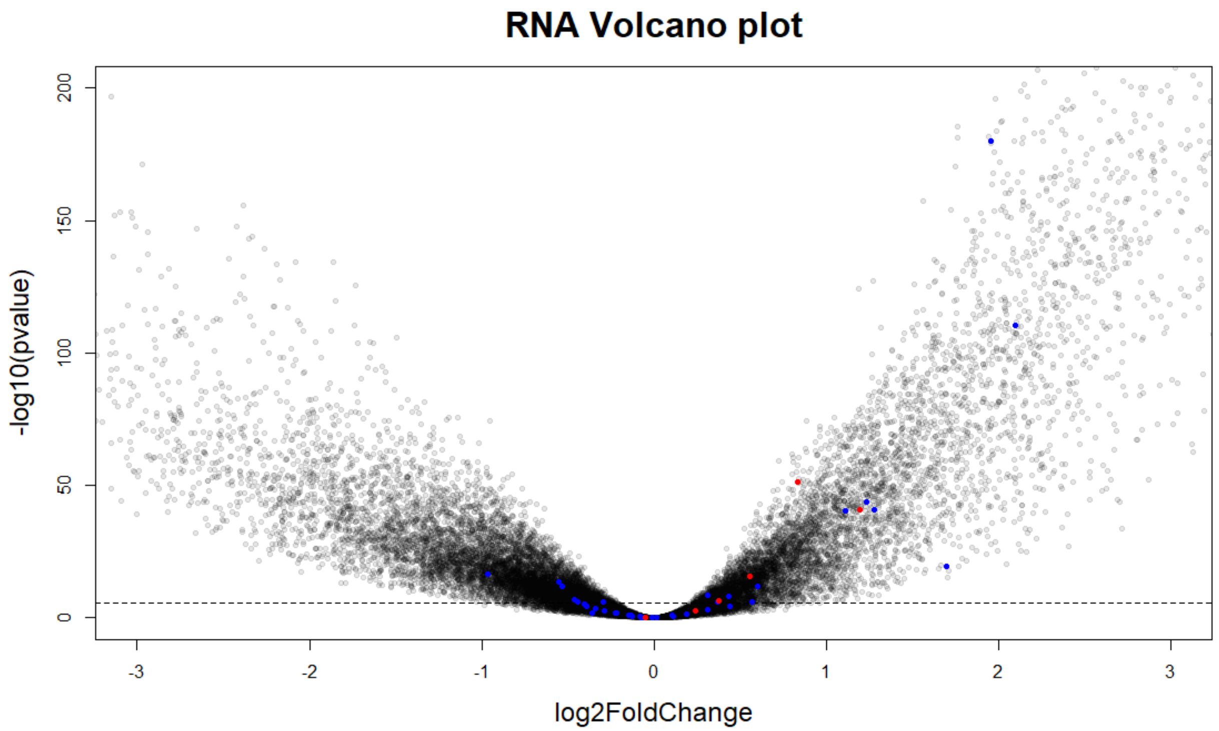

Supplemental Figure S2. Differential expression between flower and non-flower tissues. Log2Fold change of gene expression in flower tissues compared to non-flower tissues is shown (x-axis). From among the 48 relaxed genes (blue), 16 genes showed significant differential increased expression in flowers. From the 6 pollen related genes (red), 5 showed differential higher expression in flower tissues compared to non-flower tissues (blue).

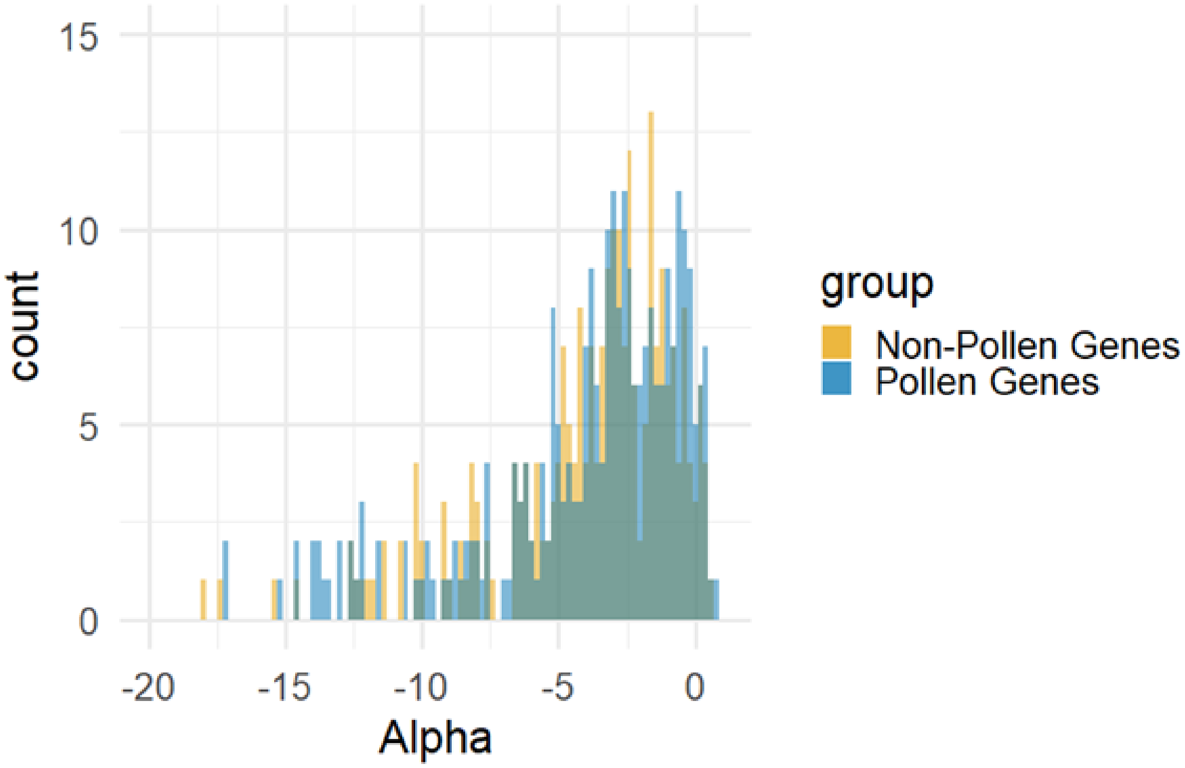

Supplemental Figure S3 Distribution of Alpha (α) value from MK test between pollen and non-pollen related genes. Histograms shown between pollen (blue) and non-pollen (orange) related genes for alpha. Non-pollen related genes are subsampled to an equal number of genes for visual comparison. No Significant difference between distributions was detected using a Chi-square test.

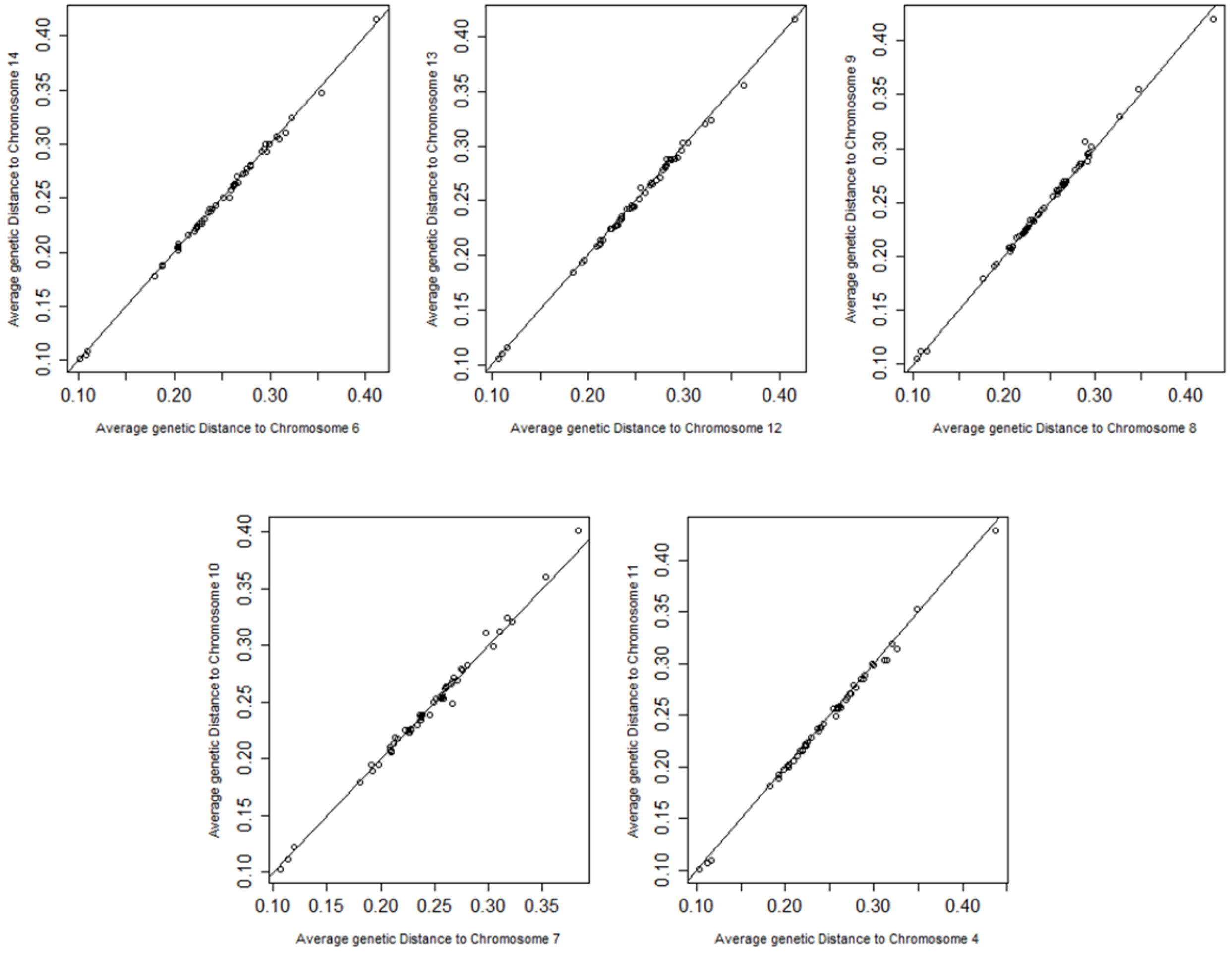

Supplemental Figure S4 Distances to each assembled genome between cassava homeologous averaged across all gens. chromosomes. Distances to each of the 52 assembled genomes were averaged over each orthologous gene. Distances were then plotted against each other, being paired by previously established ancestral homeologous chromosome pairs.

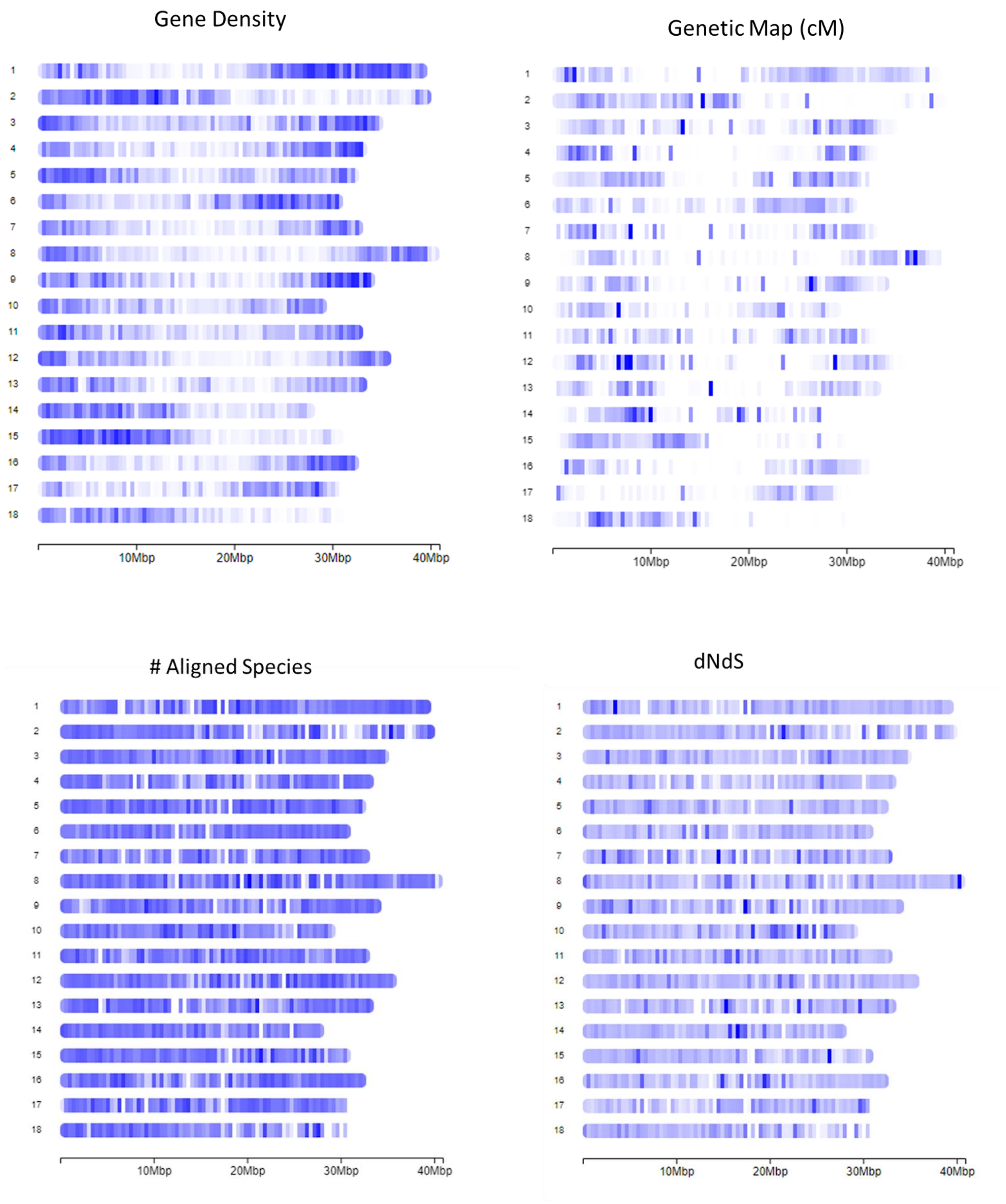

Gene Density
Genetic Map (cM)
### Aligned Species
dNdS
10Mbp
20Mbp
30Mbp
40Mbp

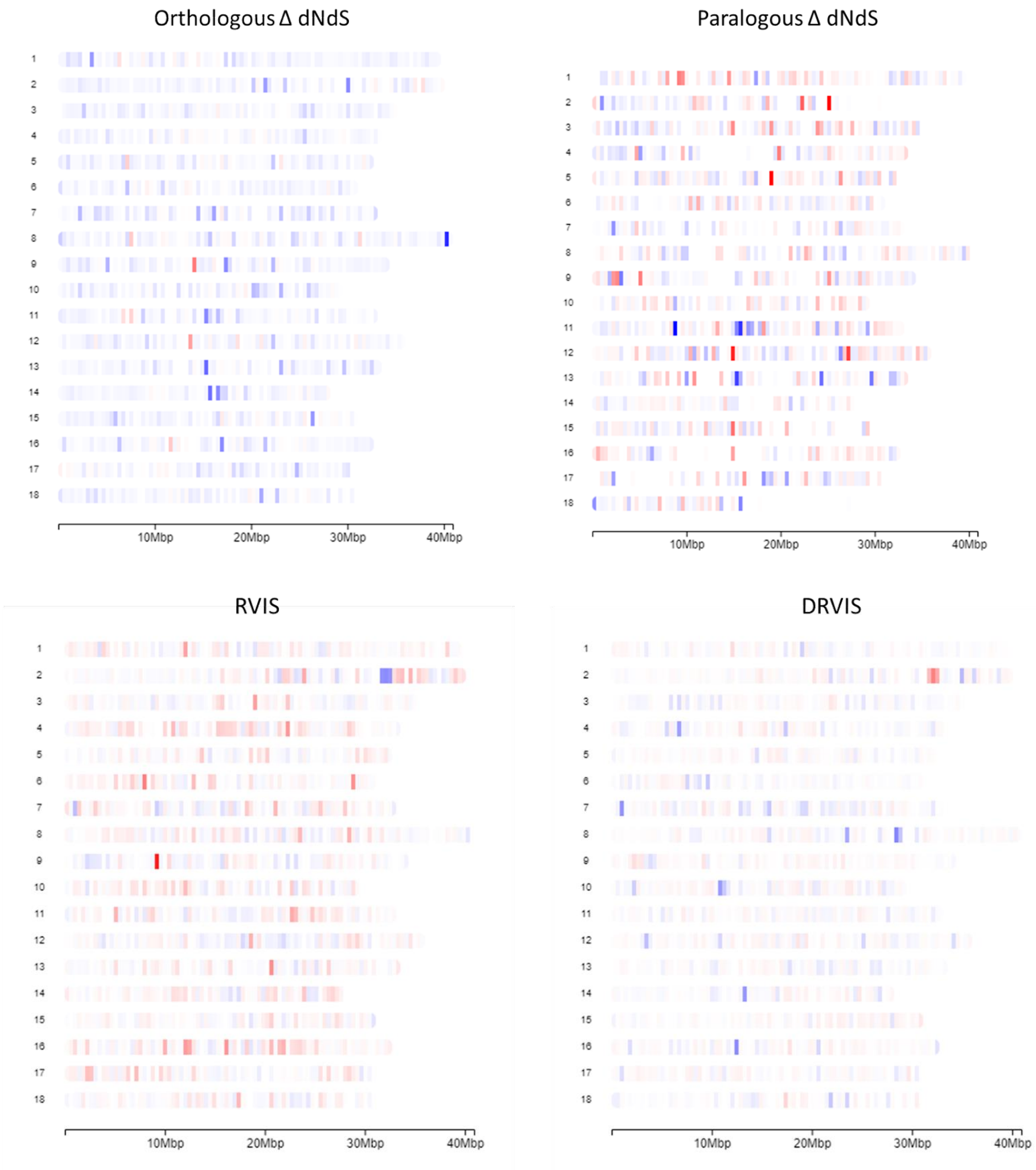

Orthologous Δ dNdS
Paralogous Δ dNdS
RVIS
DRVIS
10Mbp
20Mbp
30Mbp
40Mbp

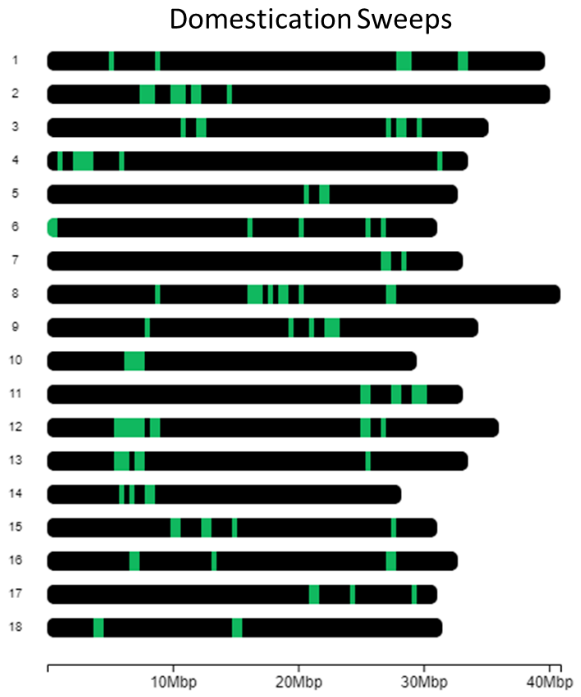

Supplemental Figure S5. Binned annotations across cassava chromosomes. Metrics including gene density, recombination rates (cM), dN/dS, number of aligned species, dN/dS differences between cassava and other species (Orthologous Δ dN/dS), dN/dS differences between gene copies (Paralogous Δ dN/dS), RVIS, DRVIS, and domestication sweeps were binned into 250kb windows for visualization.

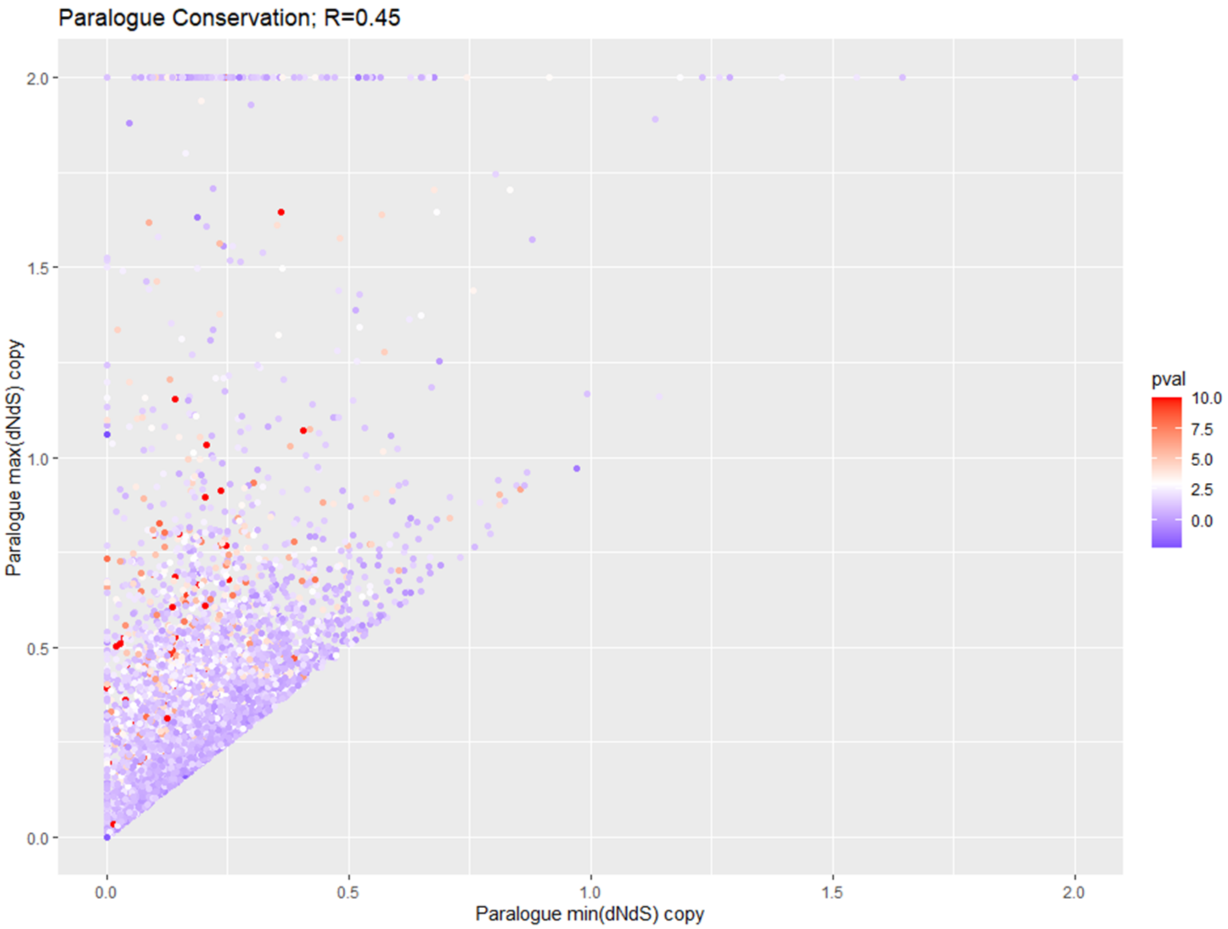

Supplemental Figure S6. Maximum and minimum values for paralogous gene copies. Among duplicated genes the dN/dS value for the most conserved copy (lowest dN/dS, x-axis) is compared to least conserved copy (highest dN/dS, y-axis). Correlation coefficient R<1 indicates asymmetrical conservation of paralogous gene copies. -log10(p-value) significance is indicated by color for differences between dN/dS in cassava and the Euphorbiaceae tree.

### Supplemental Tables

| | Genus/SRA-ID | Species/Sample-ID | Germplasm identifier | Source | Seq-type | Assembly | N50 (bp) | Raw Sequ | K-mer est | Estimated |
|---|---|---|---|---|---|---|---|---|---|---|
| 1 | Acalypha | hispida | TARS 4886 | Germplasm Resources Information Network | Short-Read | 2.16E+09 | 937 | 19.9 | NA | NA |
| 2 | Aleurites | moluccanus | PI 159043 | Germplasm Resources Information Network | Short-Read | 1.97E+09 | 1047 | 122.7 | 1550 | 79.16129 |
| 3 | Bocquillonia | castaneifolia | 20090461*A | Montgomery Botanical Garden | Short-Read | 2.97E+09 | 507 | 131.3 | 2500 | 52.52 |
| 4 | Breynia | nivosa | 957/2003*C | New York Botanic Garden | Short-Read | 2.45E+09 | 379 | 110.1 | 1910 | 57.64398 |
| 5 | Chamaesyce | celastroides | 07-0187 | US Botanic Garden | Short-Read | 3.16E+08 | 7312000 | NA | NA | NA |
| 6 | Cnidoscolus | chayamansa | 916/92*A | New York Botanic Garden | Short-Read | 2.96E+09 | 390 | 119 | 2300 | 51.73913 |
| 7 | Cnidoscolus | aconitifolius | 1996-3422-1 | Missouri Botanic Garden | Long-Read | 9.18E+08 | 1621000 | 13.36 | 505 | 26.45545 |
| 8 | Codiaeum | variegatum | 2011-1101-5 | Missouri Botanic Garden | Short-Read | 1.04E+09 | 472 | 83 | 1110 | 74.77477 |
| 9 | Croton | setigerus | W6 48715 | Germplasm Resources Information Network | Short-Read | 4.55E+09 | 879 | 42.1 | NA | NA |
| 10 | Dalechampia | spathulata | 1991-3019-1 & 78475*A | Missouri Botanic Garden & New York Botanic Garden | Short-Read | 4.74E+08 | 1508 | 29.2 | 425 | 68.70588 |
| 11 | Euphorbia | corollata | 540/2011*A | New York Botanic Garden | Short-Read | 1.75E+09 | 427 | 51.4 | 1100 | 46.72727 |
| 12 | Euphorbia | pulcherrima | 2016-0302 | US Botanic Garden | Long-Read | 1.85E+09 | 436 | 114.8 | 1680 | 68.33333 |
| 13 | Euphorbia | esula | | NCBI: ASM291907v1 | Public Assembly | 1.12E+09 | 605 | NA | NA | NA |
| 14 | Excoecaria | cochinchinensis | 2412/2004*A & 880627 | New York Botanic Garden & National Tropical Botanic Garden | Long-Read | 1.47E+09 | 62952000 | 21.64 | 1100 | 19.67273 |
| 15 | Flueggea | neowawraea | 4594/95*F | New York Botanic Garden | Short-Read | 1.4E+09 | 11818000 | 23.25 | 2490 | 9.337349 |
| 16 | Garcia | nutans | TARS 1881 | Germplasm Resources Information Network | Long-Read | 2.67E+09 | 576 | 241.7 | 1975 | 122.3797 |
| 17 | Hevea | brasiliensis | | NCBI: ASM1045892v1 | Public Assembly | 2.18E+09 | 48232000 | 22.7 | 2400 | 9.458333 |
| 18 | Hura | crepitans | 20172064 | National Tropical Botanic Garden | Short-Read | 1.47E+09 | 152719 | NA | NA | NA |
| 19 | hypericum | perforatum | | NCBI: hypericum.10X.300bp | Public Assembly | 3.3E+09 | 3453 | 31.6 | 4270 | 7.400468 |
| 20 | Jatropha | Curcas | | NCBI: RJC1_Hi-C | Public Assembly | 3.53E+08 | 37536 | NA | NA | NA |
| 21 | Jatropha | multifida | 1985-0884-1 | Missouri Botanic Garden | Short-Read | 2.67E+08 | 134284 | NA | NA | NA |
| 22 | Jatropha | podagrica | 654/70*B | New York Botanic Garden | Short-Read | 3.48E+08 | 23178 | 49.2 | 295 | 166.7797 |
| 23 | Mallotus | sp. | MIA 8875 | Germplasm Resources Information Network | Long-Read | 3.17E+08 | 23491 | 38.9 | 260 | 149.6154 |
| 24 | Mercurialis | annua | PI 279720 | Germplasm Resources Information Network | Long-Read | 1.42E+09 | 2311000 | 18.35 | 1400 | 13.10714 |
| 25 | Monadenium | arborescens | 11037*1 | Denver Botanic Garden | Short-Read | 7.3E+08 | 693576 | 11.68 | 960 | 12.16667 |
| 26 | Monadenium | elegans | 01-0829 | US Botanic Garden | Short-Read | 4.23E+09 | 828 | 38.2 | 6000 | 6.366667 |
| 27 | Monadenium | guentheri | 33013*1 | Denver Botanic Garden | Short-Read | 2.51E+09 | 700 | 19.3 | NA | NA |
| 28 | Omphalea | sp. | 1997-2689-1 | Missouri Botanic Garden | Short-Read | 4.11E+09 | 647 | 32.5 | NA | NA |
| 29 | Pedilanthus | macrocarpus | 01-1676 | US Botanic Garden | Short-Read | 1.1E+09 | 21225 | 97.9 | 1000 | 97.9 |
| 30 | Populus | deltoides | | NCBI: ASM1585260v2 | Public Assembly | 2.43E+09 | 601 | 24.8 | NA | NA |
| 31 | Populus | simonii | | NCBI: Populus_simonii_2.0 | Public Assembly | 4.29E+08 | 1838000 | NA | NA | NA |
| 32 | Populus | Trichocarpa | | NCBI: Pop_tri_v3 | Public Assembly | 4.41E+08 | 1939000 | NA | NA | NA |
| 33 | Reutealis | trisperma | PI 112679 | Germplasm Resources Information Network | Long-Read | 4.34E+08 | 552806 | NA | NA | NA |
| 34 | Rhizophora | apiculata | | NCBI: Rap_scaffold_v2 | Public Assembly | 7.23E+08 | 681198 | 23.94 | 765 | 31.29412 |
| 35 | Ricinus | Communis | | NCBI: JCVI_RCG_1.1 | Public Assembly | 2.32E+08 | 4320000 | NA | NA | NA |
| 36 | Salix | brachista | | NCBI: ASM907833v1 | Public Assembly | 3.4E+08 | 9523000 | NA | NA | NA |
| 37 | Salix | dunnii | | NCBI: SDv1.1 | Public Assembly | 3.28E+08 | 16657000 | NA | NA | NA |
| 38 | Salix | suchowensis | | NCBI: ASM1755242v1 | Public Assembly | 3.56E+08 | 2.00E+05 | NA | NA | NA |
| 39 | Salix | purpurea | | Phytozome: Salix purpurea v5.1 | Public Assembly | 1.28E+09 | 19951 | NA | NA | NA |
| 40 | Sapria | himalayana | | NCBI: Sapria_himalayana_v1 | Public Assembly | 3.29E+08 | 5083000 | NA | NA | NA |
| 41 | SRR7121446 | SAMN08770359 | | https://www.ncbi.nlm.nih.gov/pmc/articles/PMC6441391/ | Short-Read | 8.42E+08 | 3718 | 55.4665 | 679 | 81.68851 |
| 42 | SRR7121479 | SAMN08770392 | | https://www.ncbi.nlm.nih.gov/pmc/articles/PMC6441391/ | Short-Read | 8.24E+08 | 3144 | 70.1233 | 562 | 124.7746 |
| 43 | SRR7121540 | SAMN08770453 | | https://www.ncbi.nlm.nih.gov/pmc/articles/PMC6441391/ | Short-Read | 1.11E+09 | 8038 | 72.0622 | 848 | 84.97901 |
| 44 | SRR7121834 | SAMN08770747 | | https://www.ncbi.nlm.nih.gov/pmc/articles/PMC6441391/ | Short-Read | 1.14E+09 | 3634 | 61.7687 | 673 | 91.78113 |
| 45 | SRR7121842 | SAMN08770755 | | https://www.ncbi.nlm.nih.gov/pmc/articles/PMC6441391/ | Short-Read | 8.46E+08 | 2973 | 67.3669 | 554 | 121.6009 |
| 46 | SRR7121892 | SAMN08770805 | | https://www.ncbi.nlm.nih.gov/pmc/articles/PMC6441391/ | Short-Read | 1E+09 | 3482 | 67.0117 | 411 | 163.0455 |
| 47 | SRR7122023 | SAMN08770936 | | https://www.ncbi.nlm.nih.gov/pmc/articles/PMC6441391/ | Short-Read | 9.63E+08 | 5225 | 66.46 | 702 | 94.67236 |
| 48 | SRR7122040 | SAMN08770953 | | https://www.ncbi.nlm.nih.gov/pmc/articles/PMC6441391/ | Short-Read | 8.46E+08 | 2037 | 80.1765 | 1016 | 78.91388 |
| 49 | SRR7122043 | SAMN08770956 | | https://www.ncbi.nlm.nih.gov/pmc/articles/PMC6441391/ | Short-Read | 6.15E+08 | 4097 | 67.927 | 476 | 142.7038 |
| 50 | SRR7122045 | SAMN08770958 | | https://www.ncbi.nlm.nih.gov/pmc/articles/PMC6441391/ | Short-Read | 1.17E+09 | 2182 | 64.2577 | 482 | 133.3147 |
| 51 | SRR7122067 | SAMN08770980 | | https://www.ncbi.nlm.nih.gov/pmc/articles/PMC6441391/ | Short-Read | 5.49E+08 | 20286 | 65.2506 | 515 | 126.7002 |
| 52 | Stillingia | texana | W6 42922 | Germplasm Resources Information Network | Short-Read | 3.55E+09 | 693 | 35 | NA | NA |

Supplemental Table S1. Euphorbiaceae Sampling Data

| Gene | Best-hit-arabi-name | BAR eplant - AtGenExpress eFP | AtGenExpress eFP - Sexual Reproduction Related | BAR eplant - Klepikova eFP | Klepikova eFP - Sexual Reproduction Related |
|---|---|---|---|---|---|
| Manes.01G017600 | AT3G25500 | NA | No | Axis of Inflorescence | Yes |
| Manes.01G224800 | AT3G06720 | Imbibed Seed | Yes | NA | No |
| Manes.01G257400 | AT4G00500 | senescent Leaf | No | senescent Leaf | No |
| Manes.02G048560 | AT3G29090 | Dry Seed | Yes | Dry Seed | Yes |
| Manes.02G134800 | AT5G57655 | Flower Petals | Yes | Silique 1 | Yes |
| Manes.02G178800 | AT1G76850 | Cotyledons & Dry Seed | Yes | NA | No |
| Manes.02G207075 | AT3G26935 | Imbibed Seed | Yes | Stamen | Yes |
| Manes.02G213600 | AT5G58520 | 2nd Internode | No | 2nd Internode | No |
| Manes.02G218700 | AT2G26640 | Seed Stage 6 | Yes | Root Apex | No |
| Manes.02G218800 | AT2G26640 | Seed Stage 6 | Yes | Root Apex | No |
| Manes.02G222700 | AT1G79600 | Sepals | Yes | Silique 1 | Yes |
| Manes.03G130950 | AT2G32460 | Mature Pollen | Yes | Anthers | Yes |
| Manes.03G167100 | AT3G23160 | Sepals | Yes | Root | No |
| Manes.03G182600 | AT1G71830 | Seed Stage 6 | Yes | Flower | Yes |
| Manes.03G204900 | AT3G04690 | Mature Pollen | Yes | Anthers | Yes |
| Manes.04G009000 | AT5G22640 | Vegetative Rossette | No | Pedicel | Yes |
| Manes.04G017000 | AT5G60740 | Mature Pollen | Yes | Anthers | Yes |
| Manes.04G056400 | AT1G03050 | Mature Pollen | Yes | Anthers | Yes |
| Manes.04G084300 | AT4G36220 | senescent Leaf | No | senescent Leaf | No |
| Manes.04G095900 | AT5G36880 | Stamen | Yes | Stamen | Yes |
| Manes.04G153000 | AT4G24480 | Cauline Leaf | No | Cauline Leaf | No |
| Manes.04G165300 | AT5G12380 | Mature Pollen | Yes | Silique 5 | Yes |
| Manes.05G004700 | AT1G02550 | Mature Pollen | Yes | Flower 1 | Yes |
| Manes.05G041600 | AT4G00230 | Internode | No | Leaf Petiole | No |
| Manes.06G017100 | AT2G02370 | Dry Seed & Senescent Leaf | Yes | Senescent Leaf | No |
| Manes.06G064700 | AT3G18030 | Cotyledons & Dry Seed | Yes | Ovules | Yes |
| Manes.06G155400 | AT5G24090 | Stamens | Yes | Senescent Internode | No |
| Manes.08G037300 | AT2G38910 | Mature Pollen | Yes | Anthers | Yes |

| | | | | | |
|---|---|---|---|---|---|
| Manes.08G062900 | AT3G56640 | Mature Pollen | Yes | Flower 5 | Yes |
| Manes.08G109400 | AT1G56600 | Dry Seed | Yes | Dry Seed | Yes |
| Manes.09G054600 | AT1G57790 | Leaf | No | Leaf | No |
| Manes.10G036100 | AT2G01080 | Dry Seed | Yes | Young Seed | Yes |
| Manes.10G068400 | AT1G79610 | Seed Stage 7 | Yes | Silique 1 | Yes |
| Manes.10G093300 | AT5G64420 | Imbibed Seed | Yes | Germinating Seed | Yes |
| Manes.11G075500 | AT5G13930 | Petals | Yes | Anthers | Yes |
| Manes.13G101500 | AT2G22600 | Mature Pollen | Yes | Stigmatic Tissue | Yes |
| Manes.13G121200 | AT2G39980 | Dry Seed and Flower Petals | Yes | Germinating Seed and Flower Petals | Yes |
| Manes.14G087750 | AT5G60920 | Leaf | No | Root Apex | No |
| Manes.14G149800 | AT5G59720 | Petals | Yes | Internode | No |
| Manes.15G073500 | AT3G20530 | Mature Pollen | Yes | Anthers | Yes |
| Manes.15G119800 | AT2G13620 | Mature Pollen | Yes | Anthers | Yes |
| Manes.15G139100 | AT1G25240 | Mature Pollen | Yes | Anthers | Yes |
| Manes.15G146200 | AT5G42650 | Cotyledons | No | Mature Leaf | No |
| Manes.16G101200 | AT4G29750 | Vegetative Rossette | No | Leaf | No |
| Manes.17G014100 | AT2G35330 | Seed Stage 7 | Yes | Flower | Yes |
| Manes.18G019650 | AT1G27680 | Shoot Apex | No | Axis of Inflorescence | Yes |
| Manes.18G082700 | AT1G77380 | Anthers and Petals | Yes | Anthers | Yes |
| Manes.18G086900 | AT1G77120 | Cotyledons | No | Dry Seed | Yes |

Table S2. Table of tissue specific expression of Arabidopsis thaliana homologs to significantly relaxed cassava.

| GO.ID | Term | Annotated | Significant | Expected | Classic Fisher |
| --- | --- | --- | --- | --- | --- |
| GO:0006412 | translation | 656 | 160 | 64.35 | 1.1e-22 |
| GO:0016192 | vesicle-mediated transport | 651 | 111 | 63.86 | 7.2e-07 |
| GO:0006511 | ubiquitin-dependent protein catabolic pr... | 548 | 96 | 53.76 | 1.7e-06 |
| GO:0043161 | proteasome-mediated ubiquitin-dependent ... | 288 | 48 | 28.25 | 3.2e-06 |
| GO:0006499 | N-terminal protein myristoylation | 49 | 16 | 4.81 | 1.0e-05 |
| GO:0000028 | ribosomal small subunit assembly | 31 | 12 | 3.04 | 1.8e-05 |
| GO:0006886 | intracellular protein transport | 584 | 95 | 57.29 | 2.2e-05 |
| GO:0006888 | endoplasmic reticulum to Golgi vesicle-m... | 125 | 25 | 12.26 | 2.4e-05 |
| GO:0019941 | modification-dependent protein catabolic... | 565 | 104 | 55.42 | 8.0e-05 |
| GO:0048441 | petal development | 31 | 9 | 3.04 | 8.7e-05 |

Table S3. Enriched GO Terms for MK- α Genes. GO term enrichment produced from "topGO" among genes in the top 5% of MK-α scores.
